# Signal Detection of Point Targets Using Eigen-Images for Super-Resolution Ultrasound Imaging and Gas Vesicle Localization

**DOI:** 10.1101/2025.07.02.662077

**Authors:** Adree Bhattacharjee, Sydney Turner, Lu Diao, Siyuan Zhang, Sangpil Yoon

## Abstract

Accurate signal detection of ultrasound contrast agents, such as microbubble (MB) and gas vesicle (GV), in the presence of clutter and noise is essential to increase image quality in super-resolution ultrasound imaging (SRUS) and achieve precise GV localization. We developed and evaluated an eigen-image based signal detection method using singular value decomposition (SVD) and changepoint detection to automatically segment the data that are closely related to physical events such as MB flow and GV collapse. Eigen-image based method was compared with the elbow point and hard thresholding method when selecting MB signals after SVD of raw data, acquired from phantom and in vivo experiments. Image reconstructed by eigen-image based method was also compared with unregistered difference image for GV localization when moving GVs in a phantom were collapsed by ultrafast plane waves. The eigen-image based MB signal detection method resulted in higher vessel density (VD) visualization in both the phantom and in vivo mouse tumor. It also achieved increased signal-to-noise ratio (SNR) in both cases. Moreover, this method localized moving GVs more efficiently than the difference imaging method, without requiring pixel registration based on landmarks. The eigenimage based method offers a reliable and automated approach to MB and GV signal detection for both SRUS and point target localization. This approach is a valuable tool for medical imaging providing high-quality vessel images along with accurate locations of moving ultrasound contrast agents, which can be potentially translatable to clinical diagnosis and pre-clinical research.

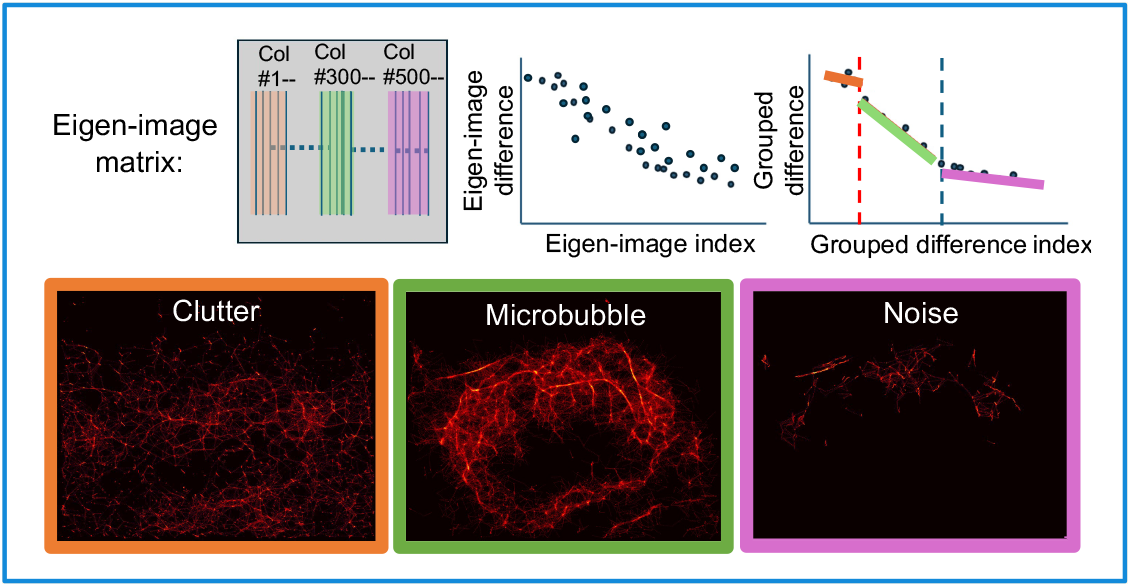

**Highlights:**

- **An approach to efficiently detect point targets including microbubbles and gas vesicles is developed using eigenimages that describe the changes of point targets over time for super-resolution ultrasound imaging and gas vesicle localization.**
- **Signals from point targets were automatically identified from clutter and noise by the changepoint detection method without prior Information.**
- **This approach enhances microvasculature visualization of tumors in mice and improves the localization of gas vesicles in a phantom.**

## I. Introduction

ULTRASOUND imaging has become an indispensable tool in biomedical research and clinical diagnosis due to its non-invasive nature, real-time imaging capabilities, and high spatial resolution [1-4]. However, traditional ultrasound imaging methods face challenges in accurately visualizing small blood vessels and capillaries, particularly when dealing with inherent noise and strong clutter signals. To enhance image quality by improving contrast, signal-to-noise ratio (SNR), and resolution, ultrasound imaging uses contrast agents such as microbubbles (MBs) and gas vesicles (GVs). MBs contain various gases that generate nonlinear acoustic responses through resonance and harmonic frequencies, which help suppress clutter in ultrasound imaging [5-12]. Recently, GVs, which are protein-based nanostructures produced by archaea and bacteria, have recently been developed as genetically encoded contrast agents for ultrasound imaging [13-17]. They offer advantages such as smaller size compared to MBs and enhanced permeability and retention (EPR) for drug delivery applications [18]. Super-resolution ultrasound (SRUS) imaging has been developed using flowing MBs in vessels and ultrafast frame rates achieved by transmitting plane waves [19-25]. For accurate selection of MB signals, it is essential to remove unwanted clutter and noise, which are common in *in vivo* ultrasound imaging. Singular value decomposition (SVD) based filtering is a key technique for achieving accurate MB localization and high-quality SRUS images [26-32].

In SVD, a sequence of two-dimensional ultrasound frames is vectorized and arranged in the columns of a measurement matrix *Y*, which is then decomposed using SVD as [26, 27, 33]:

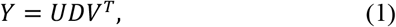

where *U* is the matrix of the left singular vectors of *Y*, each column of which is called an eigen-image, *D* is a diagonal matrix containing singular (*σ*) values, and *V^T^* is the conjugate transpose of the matrix of right singular vectors. The *σ* values in *D* are sorted in decreasing order and quantify the energy contribution of each corresponding eigen-image. The left singular vectors in *U* form a new basis that hierarchically captures the dominant spatiotemporal patterns within the data [34]. In ultrasound, clutter and tissue signals, typically slow-varying but high in amplitude, tend to occupy the first few columns in *U*. In contrast, MB or GV signals, which are more dynamic and of lower amplitude, appear in the mid-range, while noise occupies the tail end with the highest spatiotemporal frequency but lowest amplitude.

The challenge lies in determining how many of these components represent meaningful signal versus noise or clutter. SVD-based filtering has become a common approach for clutter suppression in Doppler and contrast-enhanced ultrasound imaging [26-29]. Various adaptive thresholding techniques have been proposed, including methods based on spatial singular vector similarity [28, 35], 3D clustering [29], and pixel-wise classification [36]. While these approaches provide greater flexibility, they often depend on heuristic parameters or assumptions about the data structure [37], which can lead to inconsistent performance when applied to different types of ultrasound data or imaging conditions. One widely used thresholding method is the elbow point method [38], which identifies a sharp change in the curve formed by plotting the *σ* values in descending order. This “elbow” indicates the point beyond which adding components contributes minimally to the signal and is often interpreted as the cutoff for retaining significant components. However, this method assumes a dataset with clearly separable energy levels and lacks a mathematically defined threshold, making it less reliable in complex *in vivo* conditions where clutter, MB signals, and noise may overlap. Recently, Gavish and Donoho [39] proposed a statistically grounded approach called singular value hard thresholding. Their method determines an optimal truncation threshold that minimizes the mean squared error (MSE) for low-rank approximations of matrices corrupted by Gaussian white noise and provides a reproducible threshold. However, when MB signal energy lies between clutter and noise levels, multiple rounds of thresholding may be needed to avoid signal misclassification.

To overcome these challenges, in this study, we propose to use eigen-images to extract desired signals while most approaches use the *σ* value matrix from (1). Specifically, instead of analyzing the magnitudes of *σ* values only, we analyzed changes in pixel intensities within the eigen-images using changepoint detection based on shifts in mean, variance, and slope [40, 41], to separate clutter, MB or GV signals, and noise. We compared our proposed method with the elbow point and hard thresholding method by analyzing vessel density (VD) and SNR of the reconstructed SRUS images. We validated our proposed method using a vessel-mimicking gelatin phantom with flowing MBs and an *in vivo* mouse tumor model.

To expand the utility of our eigen-image based method, we tested its ability to localize GVs as point targets in a phantom while introducing motion through lateral transducer translation. Conventional GV localization techniques, such as difference imaging [15], utilize the property of GVs collapsing above a threshold applied pressure [42]. However, this technique requires precise spatial alignment (i.e., pixel registration using a landmark) between frames acquired before and after the GV collapse event, which can be disrupted by transducer drift, physiological motion, or tissue deformation, leading to false localizations. To overcome this, we acquired high frame rate GV collapse data during continuous transducer translation using ultrafast plane wave imaging and applied our eigen-image based method without registration. We compared GV localization results from our eigen-image based method and unregistered difference imaging against ground truth using quantitative metrics including mean squared error (MSE), peak signal-to-noise ratio (PSNR), and structural similarity index (SSIM) [43, 44]. A preliminary version of this work was reported in [45]. However, the present study describes significantly different concept and approaches as we propose a new eigen-image based approach for MB signal detection and GV localization.

## II. Materials and methods

### A. Preparation of Phantoms and In Vivo Mouse Model

For the vessel-mimicking phantom experiment, a wall-less vessel-mimicking phantom, containing 8% by weight gelatin (300 Bloom, type-A, Sigma-Aldrich, Inc., St. Louis, MO) and crosslinked with a 4% paraformaldehyde, was prepared [16, 45-47]. The diameter and the length of the vessel-mimicking tunnel were 700 μm and approximately 9 cm, respectively. A second layer of phantom was prepared to mimic tissue scatterers using 8% by weight gelatin mixed with 0.1% (w/w) 40 μm silica (Si) particles (Min-U-Sil 40; Mill Creek, OH) and was placed on top of the vessel-mimicking phantom.

For the *in vivo* mouse tumor model, C57BL/6 mice (Stock: 000664, The Jackson Laboratory) were used to generate a mouse tumor model [1]. We injected approximately 0.5 million EO771 cells in phosphate buffered saline (PBS) mixed 1:1 by volume with Matrigel into the mammary fat pad. When the tumor reached 10 mm in diameter, we performed imaging.

For the GV collapse phantom, AnaGVs were isolated from *Anabaena flos-aquae* (Culture Collection of Algae and Protozoa, Scottish Marine Institute, UK), grown in G625 media containing NaNO3, KH_2_PO_4_, MgSO_4_, Na_2_SiO_3_, Na_2_CO_3_, NaHCO_3_, CaCl_2_, citric acid, EDTA, and ferric ammonium citrate. The growth process started in 15 mL tubes for 2 weeks in a 1% CO_2_ controlled incubator rotating at 100 rpm with light cycles to mimic day and night. The top layer of the solution was then added to a 1 L sterilized flask with 250 mL of the G625 growth media for an additional 3 weeks of growth in the incubator [16, 48]. GVs were concentrated using a separatory funnel, lysed with sorbitol and Solulyse, and purified by centrifugation at 350g and 4°C for 24 hours, with daily PBS replacement for 3 days. Purified GVs were stored at 4°C at OD_500_ of 20 [48] and measured by NanoDrop 2000 UV-Vis Spectrophotometer (Thermo Fisher Scientific). The well was formed by adhering a 3.2 mm diameter, 2 mm height cylinder to a gel block container lid. A 1% agarose gel with 0.01% 10 µm Si particles in Type I water at 50°C was poured into the container, and the lid rested on top. After solidifying at 4°C for 15 minutes, the lid was removed, leaving a well in the gel. A mixture of 15 µL warm gel and 5 µL GV solution was loaded into the well, solidified at 4°C for 2 minutes, and imaged with ultrasound.

### B. Ultrasound Imaging System Setup

RF data and in-phase and quadrature (IQ) data were acquired using an ultrasound imaging research platform (Vantage 256, Verasonics, Seattle, WA) connected to linear array transducers selected based on the experiment. A 3D motorized translation stage (ILS150CC, Newport Corp., Irvine, CA) was used to control the spatial positioning of the linear array transducers, allowing precise movement in lateral directions (Fig. 1a). Specifically, the L11-5v, L35-16vX, and L22-14vX transducers were used for the vessel-mimicking phantom, *in vivo* mouse tumor imaging, and GV collapse phantom experiments, respectively. These transducers have respective center frequencies of 7.6 MHz, 28 MHz, and 15.6 MHz, and each consists of a 128-element array (Fig. 1b). The transducers were connected to the Vantage 256 system (Fig. 1c), which controlled transmission sequences and acquired both RF and IQ data. Post-processing of the acquired data was performed in Matlab (Fig. 1d).

**Fig. 1.**
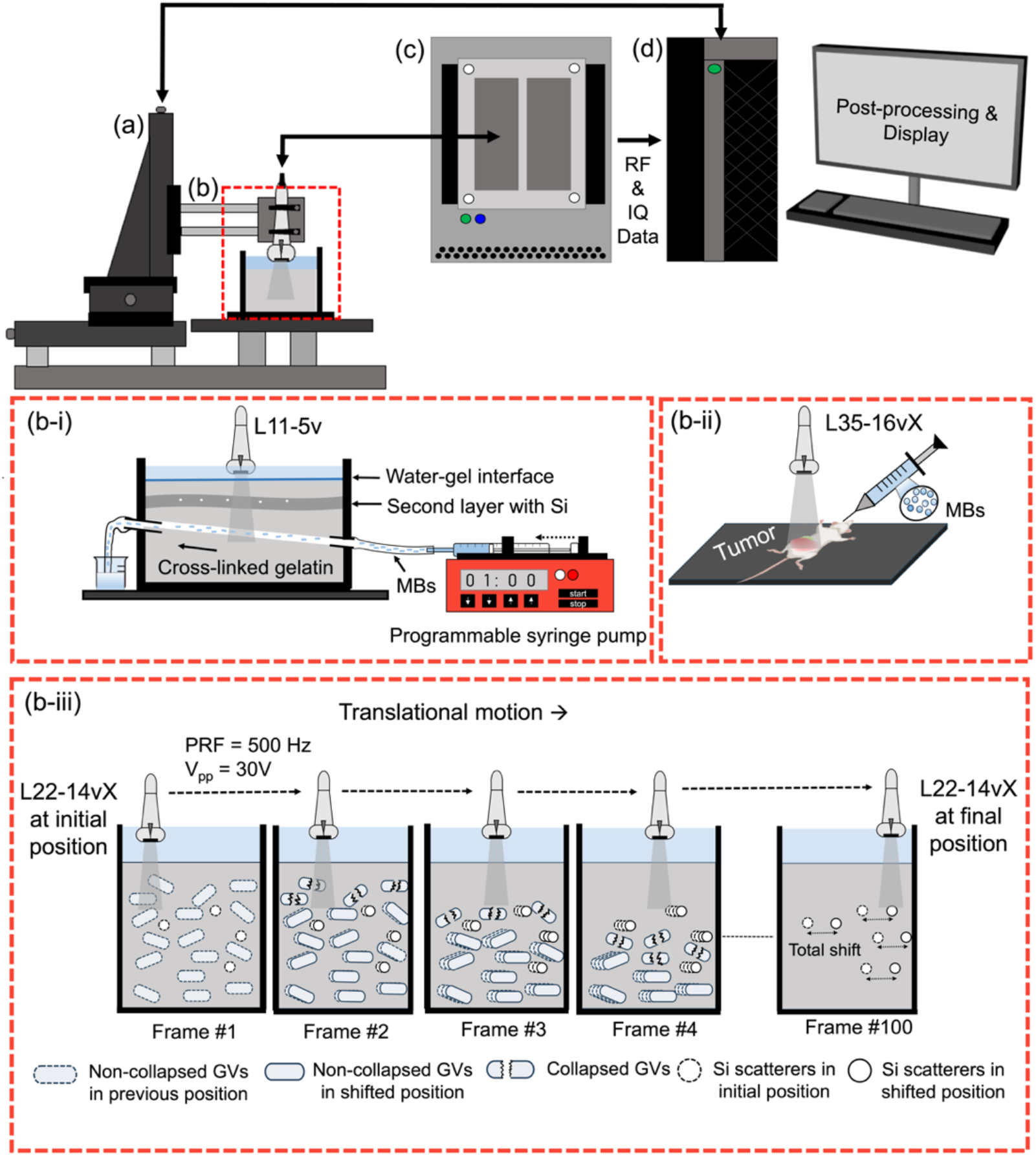
Schematic of the ultrasound imaging system setup. (a) A 3D motorized translation stage was used to control the transducer positioning. (b) A linear array transducer corresponding to specific experiment was mounted vertically above the sample to be imaged. (b-i) *Ex-vivo* vessel-mimicking phantom with a diameter of 700 μm was covered by a second layer to mimic tissue scattering. MBs were injected at a controlled flow rate into the vessel-mimicking phantom and imaged using the L11-5v transducer. (b-ii) A tumor-bearing mouse imaging setup was used to image *in vivo* tumor vessels. MBs were retro-orbitally injected for ultrasound imaging using the L35-16vX transducer. (b-iii) In the GV collapse phantom experiment, a gelatin phantom embedded with AnaGVs and Si particles was imaged while the L22-14vX transducer was translated laterally at 0.6 mm/s to simulate motion of an imaging target. Plane waves with *V_pp_* of 30 V and pulse repetition frequency (PRF) of 500 Hz were transmitted continuously to acquire GV collapse data. Frames show the initial non-collapsed GVs and Si particles in initial transducer position, collapse of the GVs, transducer motion across intermediate frames, and collapsed GVs and Si particles in shifted transducer position. (c) The Vantage 256 ultrasound imaging system was used to control transmission control and data acquisition. (d) Acquired data were post-processed using Matlab on a personal desktop computer.

### C. Imaging Procedure and Data Acquisition

For the vessel-mimicking phantom experiment, a 3 Ml syringe and a 16-gauge needle were used to administer 3 mL MBs with a concentration of 0.4 × 10^9^ MBs/mL (Lumason, Bracco Diagnostics Inc.) into the vessel-mimicking phantom at a flow rate of 1.5 mL/min. IQ data were collected using Vantage 256 connected with the L11-5v linear array transducer (Verasonics, Seattle, WA). The transducer has axial and lateral resolutions of 131 μm and 300 μm, respectively. One and a half cycle sinusoidal signal with a pulse width of 44 nanoseconds was used with a peak-to-peak voltage (*V_pp_*) of 10 V. The experiment had an effective frame rate of 500 Hz. We acquired 200 compounded IQ data sets, each consisting of five angles (ranging from −7° to 7° with 3.5° increments) over a duration of up to 400 milliseconds (Fig. 1b-i).

For *in vivo* mouse tumor imaging, we followed a similar protocol for *in vivo* mouse imaging described in our previous study [1]. A 1 mL syringe and 27-gauge needle were used to inject 150 μL (0.75 – 2.0 × 10^9^ MBs/mL) of MBs with a mean diameter of 1.5–2.5 μm through the retro-orbital of the mouse. We imaged mouse tumor using Vantage 256 with the L35-16vX linear array transducer (Verasonics, Seattle, WA). Plane waves were emitted at five different angles from −7° to 7° at 3.5° steps, with an effective frame rate of 500 Hz. One cycle sinusoidal signal was used with a *V_pp_* of 10 V. We collected 1000 compounded IQ data sets over 2 seconds, with pixel resolutions of 48.125 μm in the axial direction and 62 μm in the lateral direction (Fig. 1b-ii).

For the GV collapse phantom experiment under lateral motion, a GV-embedded gelatin phantom was imaged using the L22-14vX linear array transducer (Verasonics, Seattle, WA). The transducer was mounted on a motorized stage and translated horizontally at a constant speed of 0.6 mm/s, resulting in a lateral shift of approximately 118 μm (∼3 pixels) over the 200 milliseconds acquisition period. One and a half cycle pulses with *V_pp_* of 30 V having pulse width of 20 nanoseconds were transmitted continuously at a pulse repetition frequency (PRF) of 500 Hz to cavitate the GVs, and IQ data were acquired for 100 frames over 200 milliseconds (Fig. 1b-iii). A video capturing the real-time response of GVs to ultrasound exposure to induce GV collapse is provided in Supplementary Video.

### D. Implementation of SVD Thresholding

After performing SVD using (1), the *D* matrix was analyzed to extract MB signals for the vessel phantom and *in vivo* mouse imaging for the implementation of both the elbow point and hard thresholding methods. In contrast, the *U* matrix was used in our proposed eigen-image based method to extract MB and GV signals for vessel phantom imaging, *in vivo* mouse tumor imaging, and GV collapse phantom imaging.

#### 1) Elbow point method

The elbow point for *σ* value thresholding was determined following the approach in [45, 49]. Theoretically higher-level *σ* values correspond to clutter signals, whereas lower ones correspond to noise signal. To find desired the MB signal, we reconstructed two new filtered matrices: one using the components associated with the first set of *σ* values up to the elbow point, and another using the remaining components beyond the elbow point. These filtered matrices were obtained by multiplying the corresponding parts of the *U, D* and *V^T^* matrices. So, the filtered signals were reconstructed by:

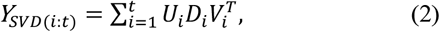

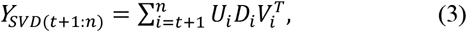

where *n* is the total number of *σ* values in *D* matrix (1) and the elbow point was found at the *t*-th threshold value. We used the filtered matrices *Y*_*SVD*(*i*:*t*)_ and *Y*_*SVD*(*t*+1:*n*)_ to reconstruct the SRUS images and observed whether the higher energy or lower energy *σ* values were associated with the MB signal.

#### 2) Hard Thresholding Method

We implemented the hard thresholding method using the source code provided by the authors in [39]. The optimal threshold location *τ_*_* was determined as described in (3) of [39]. This method retains *σ* values larger than the value at *τ_*_*, while discarding those smaller than this as noise. To ensure that MB signals were not discarded as noise, we applied the hard thresholding a second time to the discarded *σ* values from the initial step. The SRUS images were reconstructed after both rounds of thresholding, using the matrix components within the index range of the retained *σ* values each time, along with their corresponding *U* and *V^T^* matrices using (2) and (3).

#### 3) Proposed Eigen-Image Based Method Using Changepoint Detection

The proposed eigen-image based method analyzed the variations across consecutive eigen-images to identify clutter signals such as mouse movement and tissue scatterers, MB signals, and background noise. Specifically, we analyzed the *U* matrix obtained from (1) by calculating the absolute pixel intensity difference between two consecutive eigen-images. If *I_t_*(*x,y*) represents the pixel intensity at position *(x,y)* in eigenimage *t*, and *I*_*t*−1_(*x,y*) is the pixel intensity at the same position in the previous image *t*−1, *X* and *Y* are the total number of pixels along horizontal and vertical directions. Total pixel difference *E_t_* for eigen-image *t* with respect to image *t*−1 is,

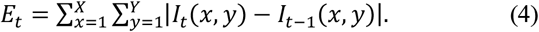

We then approximated *E_t_* for all the eigen-images grouping them into integer multiples of the total number of eigen-images in *U* matrix. For each group, we calculated the product of the mean and standard deviation to determine the mean intensity difference and variance within the group. Specifically, for a group *g* containing *M* consecutive eigen-image differences, the mean intensity difference *μ_g_* and the variance *σ_g_* were computed as:

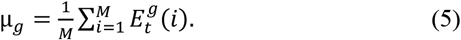

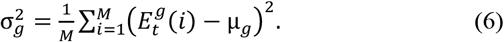

The product of the mean and standard deviation, *P_g_*, was then used to calculate the intensity variation within each group, which can be expressed as:

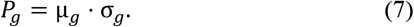

Then we applied the changepoint detection method on this calculated *P_g_* data [40]. This method identified the indices where significant changes occurred in the data with respect to pixel intensities. In this study, changepoint detection was used to observe and classify changes in the slope of *P_g_*, and to model the segments between changepoints using linear functions. To enhance the suitability of the data for analysis, a logarithmic transformation was applied before detecting change points. Matlab’s “findchangepts.m” function was employed to identify points where significant shifts occur [50]. Further details are provided in Supplementary Information Section 1 ((S1) – (S4)). If the detected changepoint index is *CP*_1_ for *M*-frame difference grouping size, then the corresponding eigen-image index *Im*_1_ is,

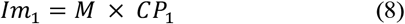

Directly applying changepoint detection to the raw frame-to-frame *E_t_* values in (4) can result in over-segmentation caused by high-frequency noise that do not correspond to meaningful changes. To mitigate this, we grouped the *E_t_* values into short, fixed-length intervals and computed their averages and variances (*P_g_* in (7)) to enhance the detection of significant structural transitions while reducing the influence of insignificant variations. However, for datasets with fewer frames or when transient behavior is expected, the changepoint method may also be directly applied to the *E_t_* values.

We then used the detected changepoints as threshold eigenimage indices in to reconstruct the 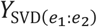 matrix for generating the SRUS images, where *e*_1_ and *e*_2_ represent the initial and final indices for each segment. This matrix can be expressed as:

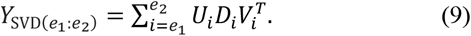

### E. SRUS Image Reconstruction and Comparison Metrics

Post-processing steps including non-local mean (NLM) filtering to enhance MB signal coherence and point spread function (PSF) characterization were performed following the methods and equation detailed in the Supplementary Information Section 2 (S5). SRUS images were reconstructed by applying MB localization using two-dimensional cross-correlation (2DCC) and L1-homotopy based compressed sensing (L1H-CS), followed by tracking of localized MB signals and accumulation of trajectories [1, 21, 51, 52].

We calculated the VD of the SRUS images obtained from the three thresholding methods. VD is defined as the ratio of the number of vessel pixels to the total number of pixels in the tumor region in the reconstructed SRUS images [1] which can be expressed as:

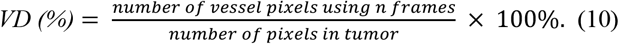

Additionally, we estimated SNR by dividing the average MB signals in the region of interest (ROI) by the standard deviation of the signal in that ROI, and is given by:

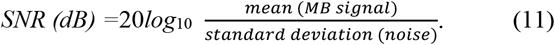

### F. GV Localization Methods and Comparison Metrics

To apply our proposed method on the GV collapse phantom data, at first, we applied SVD on the acquired dataset that captured the GV collapse event using (1). Then we applied the changepoint detection method using (4)–(7) on the *E_t_* values (4) in this case, instead of *P_g_*(7), because GV collapse is a transient event that is better characterized through the dynamics in *E_t_* values, which are the direct metrics of variation across the eigen-images. Then we reconstructed the 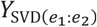 matrix in (9) using the threshold indices determined by the changepoint detection method. This method extracted GV-related signals and reconstructed the 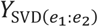 matrix suppressing the motion artifacts and noise. The first column of this matrix was selected as the GV localization image by our proposed method because it corresponded to the initial brightness mode (B-mode) frame when all GVs were still intact, but with clutter from scatterers, motion artifacts, and noise filtered out.

For comparison, a GV localization image by the unregistered difference imaging was created by subtracting a post-collapse B-mode frame from a pre-collapse B-mode frame, without pixel registration, mimicking no clear landmark was defined. To generate a ground truth image, the post-collapse frame was registered to the pre-collapse frame to compensate for motion. A bright scatterer visible throughout the sequence was used as the landmark. The lateral shift *Δx* between frames was estimated by cross-correlating intensity profiles along the lateral direction, calculated as:

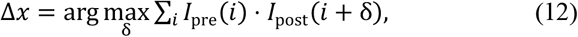

where *δ* is the variable shift being iterated over to find the optimal alignment between the two intensity profiles, *I_pre_* and *I_post_*, which are the intensity profiles from the pre-and post-collapse frames, respectively. The registered post-collapse frame was then subtracted from the pre-collapse frame to generate the ground truth GV localization image. The GV localization images obtained from our proposed method and unregistered difference imaging were compared to the registered difference image (ground truth) using MSE, PSNR, and SSIM. Detailed descriptions and equations for MSE, PSNR, and SSIM are provided in Supplementary Information Section 3 ((S6) − (S8)).

## III. RESULTS

### A. B-mode and Eigen-Image Representations

To demonstrate the advantage of eigen-images over conventional B-mode images, we illustrated some representative B-mode and eigen-images for both phantom and *in vivo* tumor datasets. Eigen-images provided clearer separation of signal components such as clutter, MB signals, and noise, compared to the superimposed appearance of these signals in B-mode images. Detailed results and representative images illustrating this comparison are provided in Supplementary Information Section 4.1 (Fig. S1).

### B. Elbow Point Method

After applying SVD to the reshaped IQ data matrix *Y* from the vessel-mimicking phantom experiment, the resulting *D* matrix was a 200 × 200 diagonal matrix containing 200 *σ* values, corresponding to the 200 frames in the original dataset. The normalized, log-scaled *σ* values were plotted (Fig. 2a). Using the elbow point method, the 16th *σ* value was identified as the threshold separating higher energy (1–16th) *σ* values (Fig. 2a, green points) from lower energy (17–200th) *σ* values (Fig. 2a, red points). Reconstructing the SRUS image using *Y*_*SVD*(1:16)_ matrix (*i* = 1 to *t* = 16 in (2)) revealed clutter signals resembling the tissue-mimicking gel background (Fig. 2b). In contrast, the lower energy *σ* values reflected MB signals and noise, producing a distinct vessel structure in the SRUS image using *Y*_*SVD*(17:200)_ matrix (*t* = 16 to *n* = 200 in (3)) (Fig. 2c). Similarly, SVD was applied to the reshaped *in vivo* mouse tumor IQ data containing 1000 frames, resulting in a *D* matrix with 1000 *σ* values (Fig. 2d). The elbow point method identified the 163rd *σ* value as the transitional point (blue point), marking 1–163rd values as higher energy *σ* values (Fig. 2d, green points) and 164–1000th as lower energy *σ* values (Fig. 2d, red points). The SRUS image reconstructed from *Y*_*SVD*(1:163)_ matrix in (2) appeared dominated by tissue clutter (Fig. 2e), while the lower energy *σ* values revealed MB signals and noise, resulting in a clear depiction of tumor vasculature in the SRUS image using *Y*_*SVD*(164:1000)_ matrix from (3) (Fig. 2f).

**Fig. 2.**
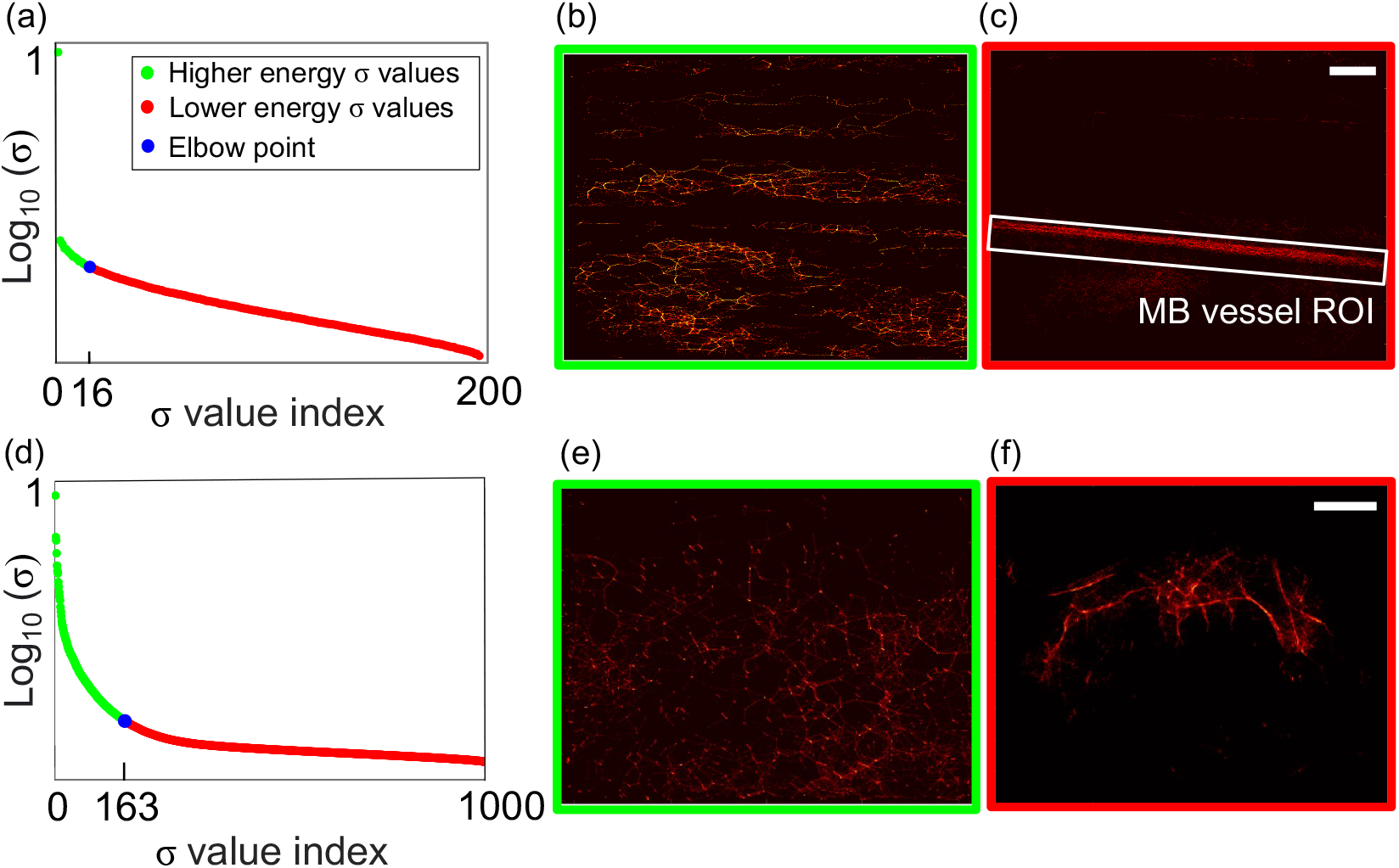
Elbow point method was applied to the vessel and *in vivo* mouse tumor data. (a) For the phantom data, elbow point method detected the 16^th^ *σ* value as a threshold (blue point) between higher energy (green points) and lower energy (red points) *σ* values from the *σ* value plot. (b) SRUS image reconstructed using *Y*_*SVD*(1:16)_matrix from (5) showed clutter signal. (c) SRUS image reconstructed using *Y*_*SVD*(17:200)_matrix from (6) represented MB signal in the vessel. Reconstructed vessel structure ROI is shown in white box. (d) For the *in vivo* data, elbow point method detected the 163^rd^ *σ* value as the threshold for higher energy and lower energy *σ* values. (e) SRUS image reconstructed using *Y*_*SVD*(1:163)_ from (5) showed tissue clutter. (f) SRUS image reconstructed using *Y*_*SVD*(164:1000)_ from (6) showed the vasculature of the tumor. White scale bars: (c) 1 mm, (f) = 10 mm.

### C. Hard Thresholding Method

Hard thresholding was applied to the decomposed *D* matrix using (3) of [39]. The first threshold retained the top 45 *σ* values with the highest energy (Fig. 3a, green points). The SRUS image reconstructed using *Y*_*SVD*(1:45)_ matrix from (2) highlighted clutter signals originating from the gelatin layer embedded with Si particles (Fig. 3b). To extract MB signals, a second hard thresholding was performed on the remaining lower-energy *σ* values (46–200th), retaining the next 35 *σ* values (Fig. 3a, red points). The SRUS image reconstructed using *Y*_*SVD*(46:80)_ matrix clearly revealed vessel structures formed by MB signals with minimal clutter (Fig. 3c). A similar hard thresholding approach was applied to the decomposed *D* matrix of the *in vivo* mouse tumor data. At first, the top 211 *σ* values were retained as higher energy components (Fig. 3d, green points). The SRUS image reconstructed using *Y*_*SVD*(1:211)_ matrix from (2) showed dominant clutter signals from tissues surrounding the tumor (Fig. 3e). To isolate MB signals, a second hard thresholding step retained the next 35 *σ* values (212–246th), representing lower energy components (Fig. 3a, red points). The SRUS image reconstructed using *Y*_*SVD*(212:246)_ matrix revealed tumor vasculature with minimal clutter (Fig. 3f).

**Fig. 3.**
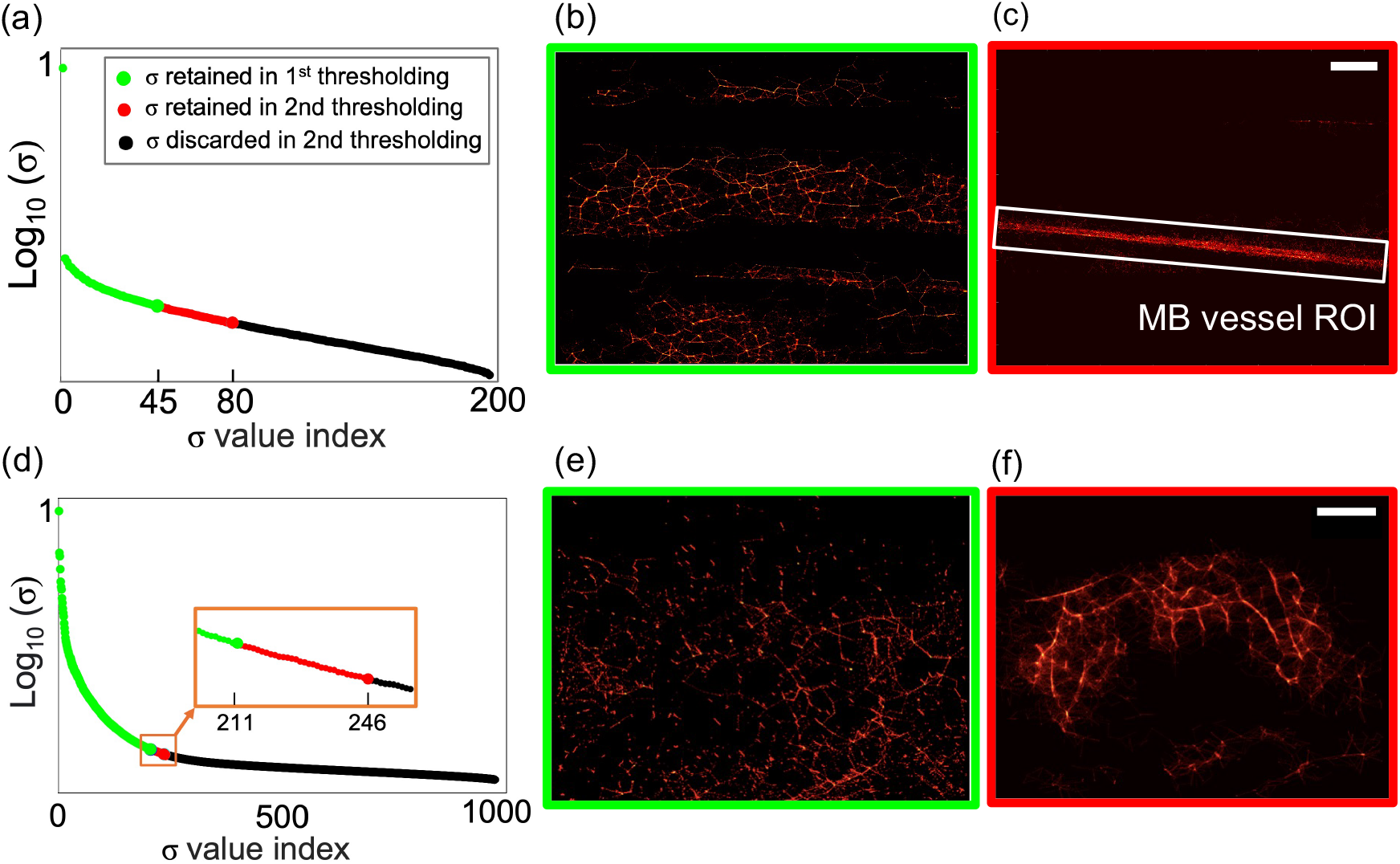
Hard thresholding method was applied to the vessel phantom and *in vivo* mouse tumor data. (a) In the *σ* value plot of the vessel phantom data, the first and second threshold points identified the 45^th^ and 80^th^ *σ* values as separations between clutter, MB signal components and noise (green, red points and black points, respectively). (b) SRUS image was reconstructed using *Y*_*SVD*(1:45)_, obtained after the first round of thresholding, which primarily corresponded to clutter signals. (c) SRUS image was reconstructed using *Y*_*SVD*(46:80)_, obtained after the second round of thresholding, representing MB signal. The remaining components were discarded as noise. Reconstructed vessel structure ROI is shown in white box. (d) For the *in vivo* data, the first and second threshold points identified the 211^st^ and 246^th^ *σ* values as separations between clutter, MB signal components and noise. (b) SRUS image was reconstructed using *Y*_*SVD*(1:211)_identified after first round of thresholding expressed clutter signal. (c) SRUS image was reconstructed using *Y*_*SVD*(212:246)_identified after the second round of thresholding expressed tumor vasculature. The remaining components were discarded as noise. White scale bars: (c) 1 mm, (f) = 10 mm.

### D. Proposed Eigen-Image Based Method

The proposed eigen-image based method was applied to both the vessel phantom and *in vivo* datasets following the procedure in Section II-D-3. At first, *E_t_* values were calculated using (4) (Figs. 4a, 5a). Then, changepoint detection applied to the *P_g_* plots consistently determined two changepoints for both datasets, segmenting the data into three parts. For the phantom data, *M* = 4 and 5 were used in (5) and (6), with changepoints detected near the 15th and 110th indices (Table I, Fig. 4b,c). For the *in vivo* tumor data, group sizes *M* = 4, 5, 10, and 20 were applied, obtaining changepoints near the 100th and 300th indices (Table II, Fig. 5b–e). In both cases, SRUS images reconstructed by using the components within the starting and ending indices of the middle segment using (9) provided detailed visualization of MB vessel structures or tumor vasculature with reduced background clutter (Figs. 4d–f, 5f–h).

**TABLE I.** OPTIMIZED CHANGEPOINTS FOR DIFFERENT *M* FOR VESSEL PHANTOM DATA USING (8)

| $M$ | $CP_1$ | $Im_1$ | $CP_2$ | $Im_2$ |
| --- | --- | --- | --- | --- |
| 4 | 4 | 16 | 27 | 108 |
| 5 | 3 | 15 | 22 | 110 |

**TABLE II.**
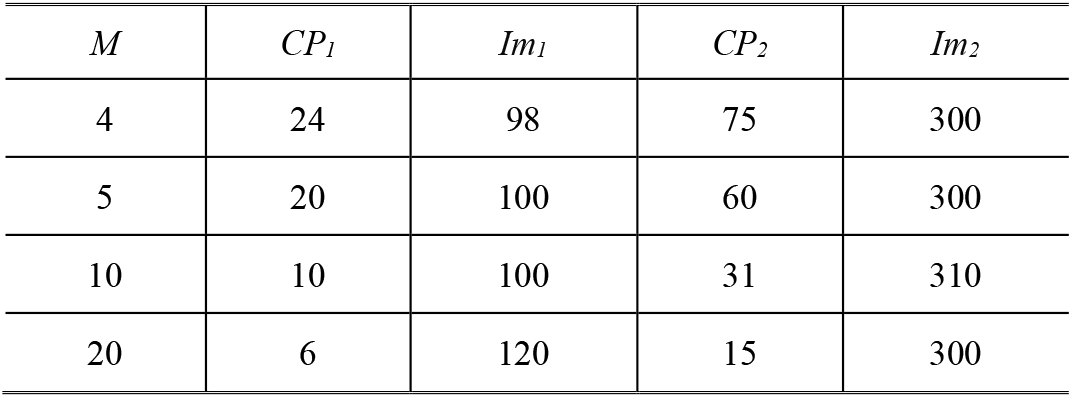
OPTIMIZED CHANGEPOINTS FOR DIFFERENT *M* FOR *IN VIVO* MOUSE TUMOR DATA USING (8)

**Fig. 4.**
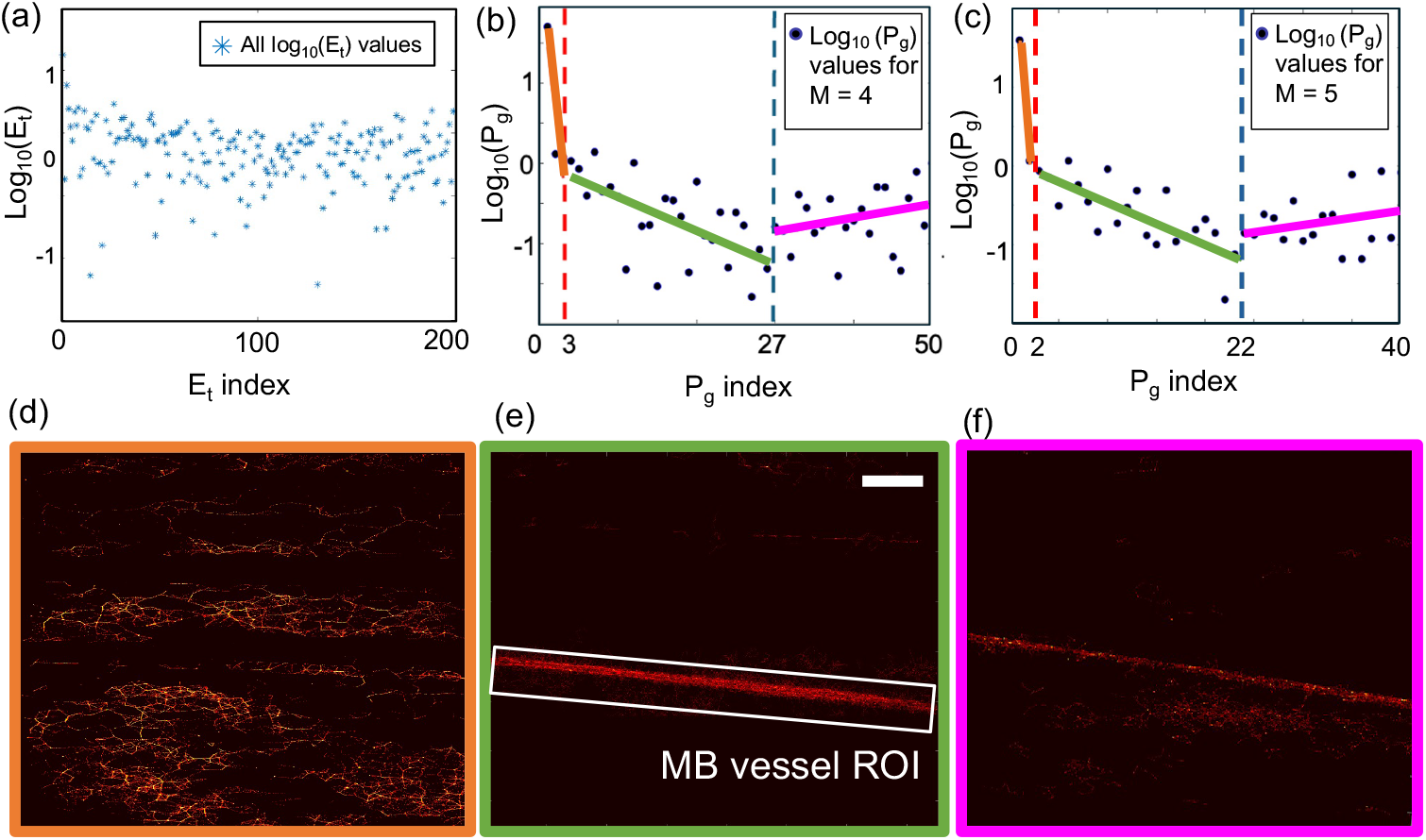
The eigen-image based method was applied to the vessel phantom data. (a) Consecutive eigen-image absolute pixel intensity differences (*E_t_*) were calculated and plotted using (4) from the *U* matrix of the vessel phantom data. (b),(c) Changepoint detection was applied to the product of mean and standard deviation, *P_g_* by (5)–(7) calculated from grouped *E_t_* values using *M* = 4 and 5 from (5) and (6), respectively. The method identified three segments with two changepoints located around the 15^th^ and 110^th^ indices represented by red and blue dotted lines. (d) SRUS images were reconstructed using (9) for the following index ranges: (d) *e_1_*= 1 to *e_2_*= 15 (orange dashed line), (e) *e_1_*= 16 to *e_2_*= 110 (green dashed line), and (f) *e_1_*= 111 to *e_2_*= 200 (pink dashed line). The second range, identified by the changepoints, produced the clearest visualization of the MB vessels. White scale bar = 10 mm.

**Fig. 5.**
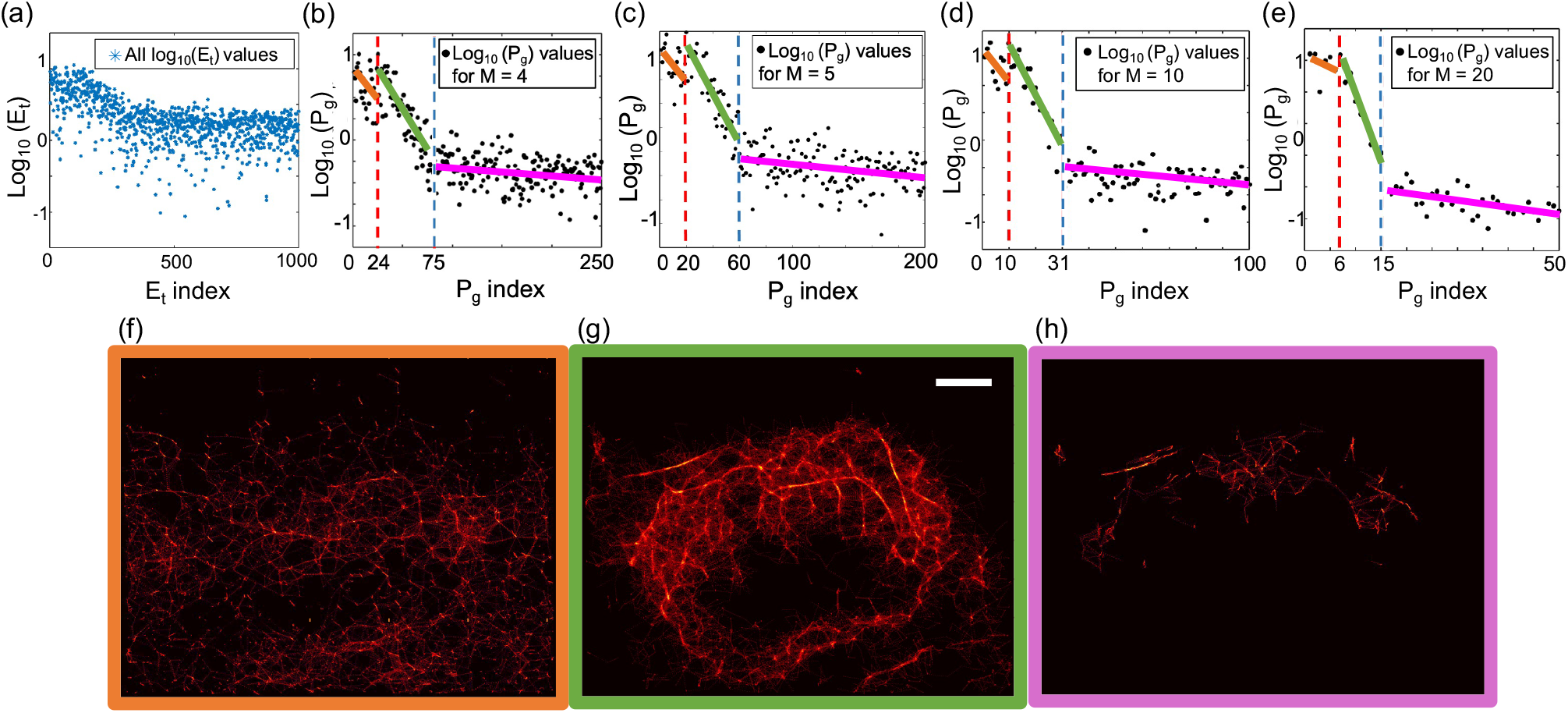
The eigen-image method was applied to *in vivo* mouse tumor data. (a) Consecutive eigen-image intensity differences (*E_t_*) were calculated from the *U* matrix. (b,e) Changepoint detection was performed on grouped *E_t_* statistics (*P_g_*) using group sizes *M* = 4, 5, 10, and 20, identifying two changepoints near indices 100 and 300 (red and blue dotted lines). (d) SRUS images were reconstructed for (f) indices 1–100 (orange dashed line), (g) 101–300 (green dashed line), and (h) 301–1000 (pink dashed line). The 101–300 range provided the clearest tumor vasculature. White scale bar = 1 mm.

### E. Comparison of 2DCC SRUS images by three methods

We compared the eigen-image, elbow point, and hard thresholding methods for SRUS imaging using the vessel phantom and *in vivo* tumor datasets. Representative 2DCC SRUS images of the vessel phantom reconstructed with 200 frames are shown in Fig. 6a–c for all three methods. The VD and SNR of these SRUS images, calculated using (10) and (11), are plotted in Fig. 6d,e. The eigen-image based method consistently produced higher VD and SNR, especially at higher frame counts, reaching 76% VD and 8.40 dB SNR at 200 frames. While the elbow point method showed performance close to that of the eigen-image based method, the hard thresholding method consistently lagged.

**Fig. 6.**
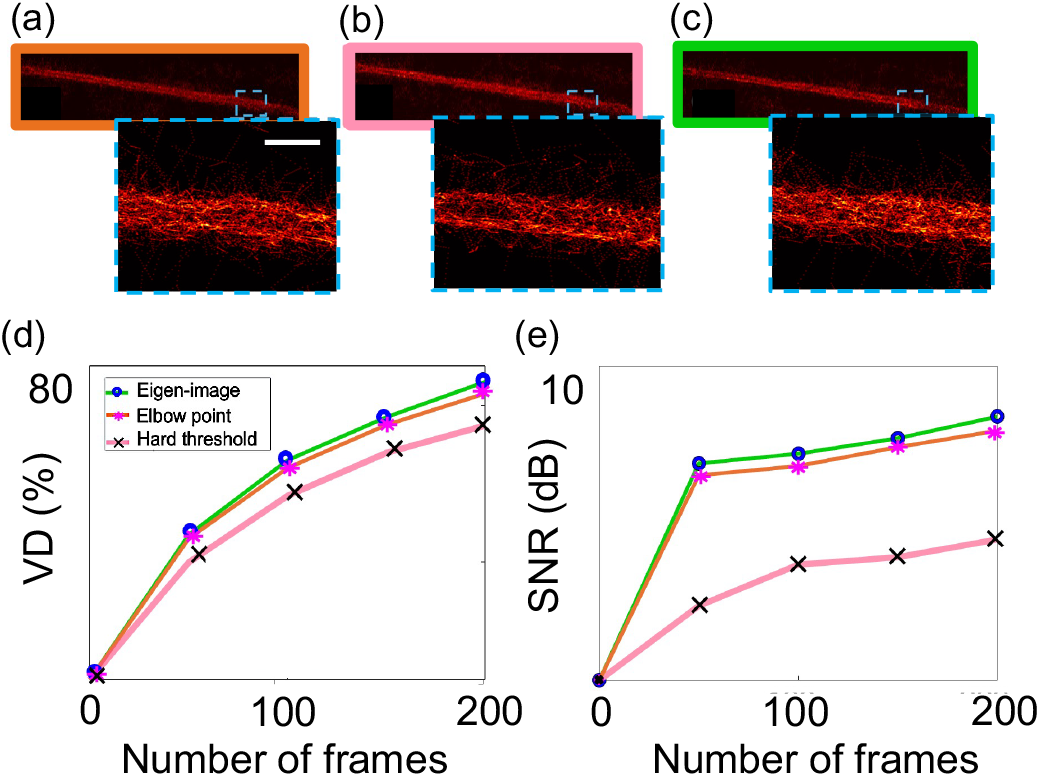
Comparison graphs of (a) VD and (b) SNR in vessel phantom imaging were plotted for different number of frames taken for generating the SRUS image for each method. SRUS images were reconstructed with 200 frames from (c) elbow point method (d) hard thresholding method (e) eigen-image based methods. White scale bar = 1 mm.

For the *in vivo* mouse tumor dataset, representative 2DCC SRUS images reconstructed with 1000 frames are shown in Fig. 7a–c, with corresponding B-mode images overlaid with SRUS images and tumor boundaries depicted in Fig. 7d–f. The VD and SNR of these SRUS images are plotted in Fig. 7g,h. The superiority of the eigen-image based method became even more evident with increasing frame counts. At 1000 frames, the eigen-image based method achieved 37.97% VD and 8.20 dB SNR, whereas the elbow point and hard thresholding methods reached only around 15–18% VD, with SNR values up to 7 dB. This performance trend was consistently observed across all other tested frame counts (50, 100, 200, and 500). It was observed that, for the *in vivo* case, the hard thresholding method performed better than the elbow point method.

**Fig. 7.**
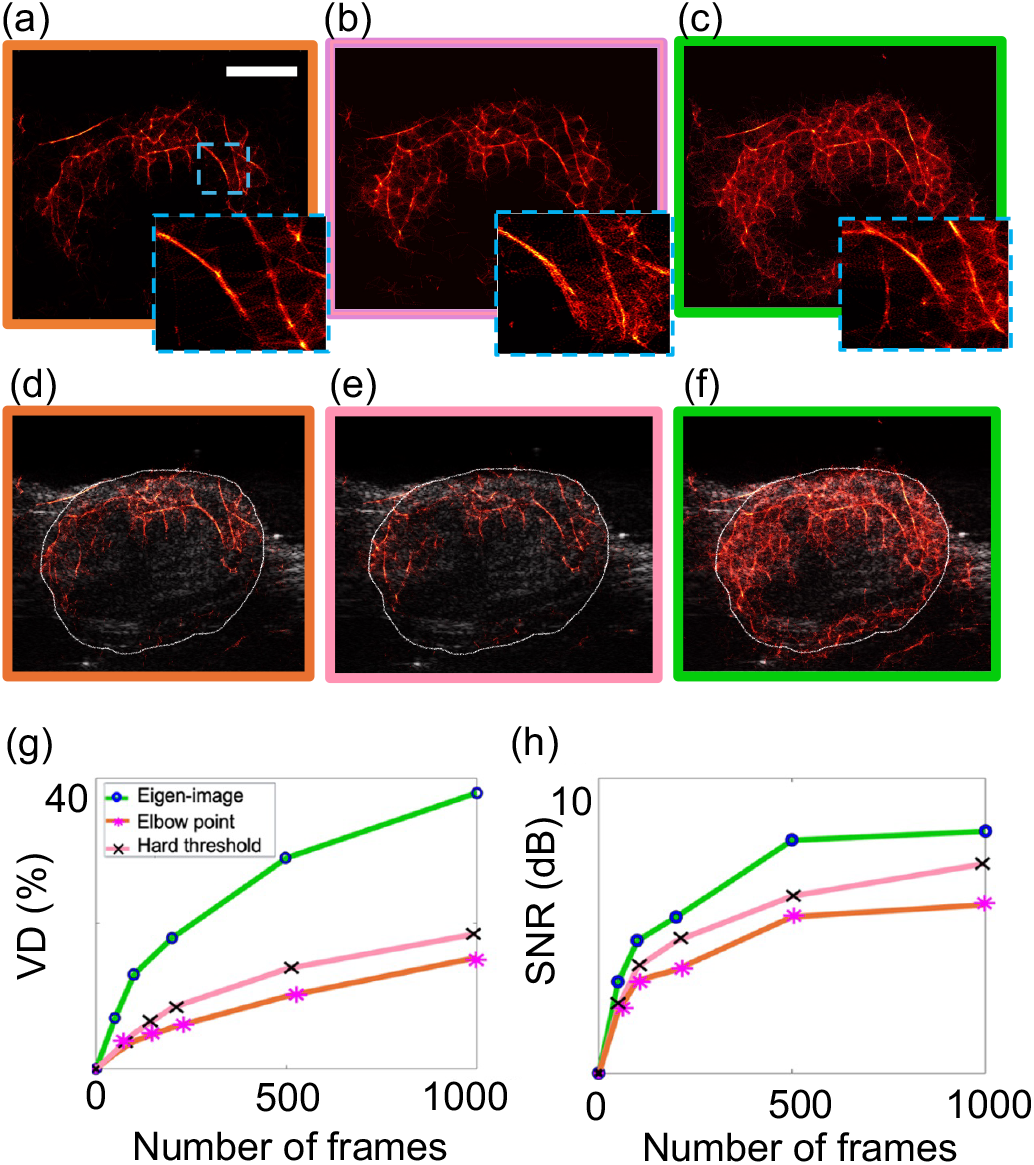
Comparison of (a) VD and (b) SNR in *in vivo* imaging for different frame counts used in SRUS reconstruction. SRUS images with 1000 frames using (c) elbow-point, (d) hard thresholding, and (e) eigen-image methods. B-mode images overlaid with SRUS and tumor boundary (white outline) for (f) elbow-point, (g) hard thresholding, and (h) eigenimage methods. White scale bar = 1 mm.

### F. L1H-CS SRUS Images

To further validate the consistency of the proposed method, VD and SNR comparisons were performed using L1H-CS localization based SRUS images [1]. As summarized in Supplementary Information Section 4.2 (Fig. S2), the eigenimage based method consistently produced higher VD and SNR for L1H-CS localization as well, compared to the elbow point and hard thresholding approaches for both phantom and *in vivo* datasets.

### G. GV Localization by Eigen-Image Based Method

We explored the proposed eigen-image based method as a potential approach to localize GVs without pixel registration. Fig. S3 in Supplementary Information Section 4.3 shows the shift of a stationary scatterer caused by transducer motion during the GV collapse event and the spatial registration used to correct this shift. The 100-frame data from this experiment were analyzed following the procedure in Section II-F (Fig. 8). This analysis identified two changepoints at indices 3 and 61, segmenting the data into three regions corresponding to clutter, GV collapse activity, and noise (Fig. 8a). The first segment (*E_t_* indices 1–2 in Fig. 8a) captured background clutter, consistent with eigen-images 1 and 2 (Fig. 8b,c). The second segment (indices 3–61) corresponded to GV collapse activity. Eigenimages 9 and 32 (Fig. 8d,e) show GV signals at various depths that correlated to GV collapse. The third segment (indices 62– 100) represents noise, as seen in eigen-images 90 and 100 (Fig. 8f,g). To compare the proposed method with conventional difference imaging, a ground truth image was generated by subtracting the 100th B-mode frame from the 1st B-mode frame after spatial registration (Fig. 9a). The GV localization image from the unregistered difference image (Fig. 9b) showed two bright spots within the ROI due to lateral motion, resulting in false positive signals unrelated to actual GV. In contrast, the first column of the reconstructed *Y*_SVD(3:61)_ matrix obtained by the proposed eigen-image based method produced a GV localization image showing a single bright spot (Fig. 9c), closely matching the ground truth. Quantitative comparison using (S6)–(S8) of the Supplementary Information Section 3 demonstrated lower MSE and higher PSNR for the eigen-image based method relative to the unregistered difference imaging approach, shown in Table III.

**TABLE III.** QUANTITATIVE COMPARISON OF GV LOCALIZATION IMAGES EVALUATED AGAINST THE GROUND TRUTH IN FIG. 9(a-c)

| Comparison parameters | Eigen-image based | Unregistered difference image based |
| --- | --- | --- |
| MSE | 0.0203 | 0.0321 |
| PSNR | 17 dB | 15 dB |
| SSIM | 0.8 | 0.8 |

**Fig. 8.**
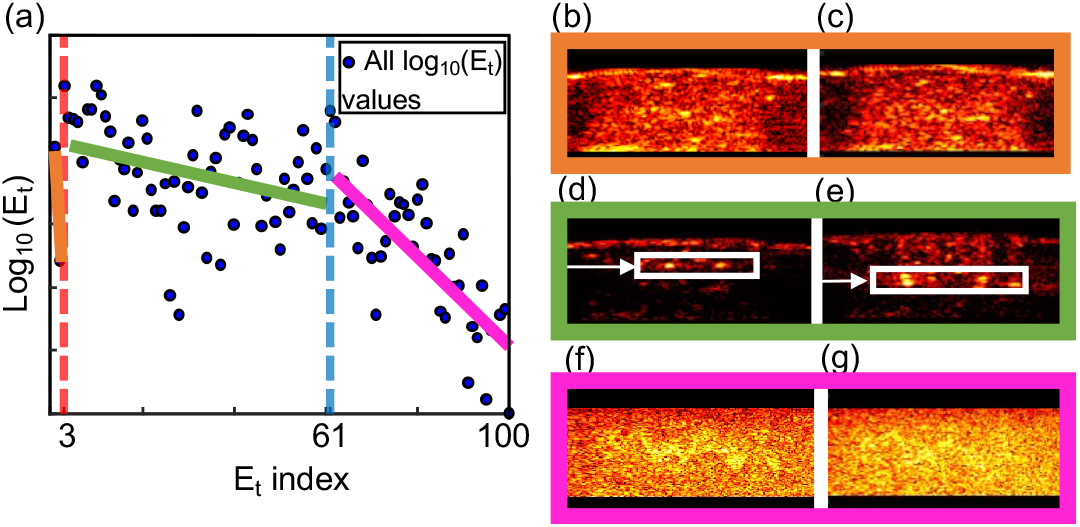
(a) Eigen-image based method using changepoint detection segmented the GV collapse data into three segments based on changepoint detection: first segment consisting of indices 1–2 (clutter), second segment consisting of indices 3–61 (GV collapse activity), and third one consisting of indices 62–100 (noise), effectively isolating the GV collapse event. The first changepoint (red dotted vertical line) separates clutter from GV activity at *E_t_* of 3, while the second changepoint (blue dotted vertical line) marks the transition from GV activity to noise at *E_t_* of 61. (b),(c) Clutter representative eigen-images 1 and 2 captured background scatterer motion. (d),(e) Eigen-images 9 and 32 showing GV collapse at increasing depth, indicating successive cavitation. White boxes are highlighting the GV region. (f),(g) Eigenimages 90 and 100 from the third segment are predominantly representing noise signal.

**Fig. 9.**
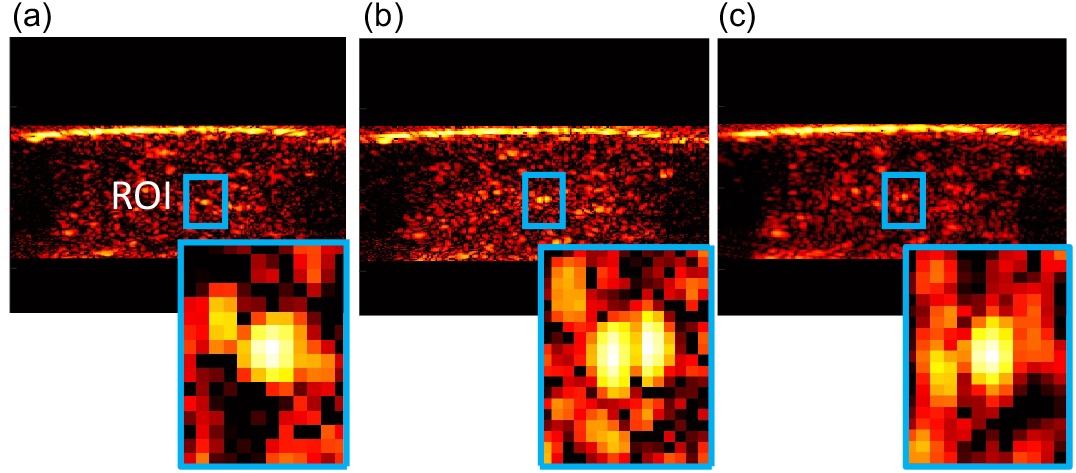
(a) Ground truth image of GV localization was generated by subtracting registered B-mode frame 100 from frame 1. (b) GV localization image from the unregistered difference imaging between B-mode frames 100 and 1 showed two dots. (c) GV localization image from our eigen-image based method showed one bright spot, similar to the ground truth. Insets show zoomed ROI comparisons across all three methods

## IV.Discussion

In this study, we introduce an eigen-image based method that uses eigen-images obtained from the SVD of the input dataset to detect and localize signals from point targets such as MBs and GVs. Eigen-images were automatically categorized based on the differences in pixel intensities between consecutive eigen-images using changepoint detection. We implemented this method to extract MB signals from a vessel mimicking phantom data and *in vivo* mouse tumor data and compared the performance of this method with the elbow point and hard thresholding methods. We extended this approach to GV localization in a phantom by tracking GV collapsing. Moving MBs in vessels and collapsing GVs were efficiently tracked and localized by proposed eigen-image based method, utilizing the fact that moving MBs and collapsing GVs present similar effects to enhance contrast and precise localization of point targets that are smaller than the diffraction limit.

The elbow point method does not efficiently categorize signals with different physical meaning including clutter, MB, and noise. It is easy to implement but does not handle complex data well, particularly in datasets with a higher level of noise clutter, specifically in *in vivo* imaging. The reason why the elbow point method performed better than the hard threshold method in a vessel-mimicking phantom is that the controlled environment of the phantom, which had less noise and fewer complexities, allowed the elbow point method to identify MB signals more effectively.

In the eigen-image method, the *U* matrix data were segmented using the changepoint method for an automated detection of different physical events. By applying changepoint detection method on the obtained values from (4)−(7), this approach automatically segmented the data according to the difference in its inherent spatial patterns, allowing precise segmentation thresholds for accurate MB signal extraction. As shown in Figs. 4 and 5, the method segmented both the phantom and *in vivo* data into three parts, which can be justified as both datasets contain strong clutter signals from tissue in phantom (Fig. 4d) or tissue and breathing motion in *in vivo* (Fig. 5f), MB signal (Figs. 4e, 5g) and noise (Figs. 4f, 5h). Interestingly, from Figs. 4 and 5, eigen-images corresponding to the clutter signal were less in number in the phantom data than in the *in vivo* data, which is reasonable considering that the phantom data did not include mouse breathing motion.

The rationale for applying the changepoint detection method to the product of the mean and variance (*P_g_* in (7)) of a group of consecutive eigen-image differences (e.g., *M* = 4, 5, 10, 20 etc. in (5) and (6)) was that it balances both intensity and variability of the consecutive eigen-images. Clutter, which was associated with stationary tissue and breathing motion, tended to have both high intensity and high variability, resulting in higher *P_g_* values. MB signals had moderate intensity and variability, leading to intermediate values, while noise, characterized by low intensity and variability, resulted in the lowest values. This combined approach improved the ability to distinguish between these categories using changepoint detection method with enhanced sensitivity and accuracy by detecting significant shifts in the statistical properties of the data.

Starting with the phantom data, the eigen-image based method demonstrated consistent superior performance, with VD values starting at 37% for 50 frames and increasing to 76% steadily with 200 frames (Fig. 6d). This steady improvement highlights the robustness of the proposed method in isolating MB signals from clutter and noise, leading to the clearest representation of vessel phantom structure. Our eigen-image based method also had the best SNR output for across all no. of frames (Fig. 6e). In the *in vivo* experiment as well, the eigenimage method outperformed both the elbow point and hard thresholding methods (Fig. 7g,h). Optimized changepoint indices remained nearly uniform across different frame group sizes as seen in Tables (I) and (II), further confirming the reliability of this approach in controlled environments. One important aspect to discuss is the grouping size when using Larger grouping sizes are more suitable for datasets with more frames (e.g., we tested *M* = 4, 5, 10, and 20 for a 1000-frame *in vivo* dataset using (5) and (6)), as it reduces sensitivity to minor fluctuations and improves the stability of trend detection.

To further validate the robustness of our eigen-image based method, we evaluated it for L1H-CS localization based SRUS images, along with 2DCC, assessing the performance consistency by VD and SNR across different localization techniques. The results in Fig. S2 in Supplementary Information Section 4.2 confirmed the consistency in performance across different reconstruction algorithms. L1H-CS consistently shows higher VD and SNR compared to 2DCC regardless of thresholding methods, which correlates with the results in our previous work [1].

We further evaluated the eigen-image based method for GV localization using a laterally moving phantom by translating the linear array transducer to create apparent GV motion. The registered difference image was the ground truth for comparison with unregistered difference imaging and our method. The reason for this comparative study was to find an alternative to difference imaging with pixel registration method, since in practice, precise landmark for registration is hard to be estimated because of biological motion, tissue deformation, and transducer drift in *in vivo* imaging. Our proposed eigen-image based method addresses this limitation by analyzing the *U* matrix using changepoint detection method to segment rigid body movement (eigen-images 1−2), GV collapse event (3−61), and noise (62−100) linked to physical events (Fig. 8). Using (9), we reconstructed *Y*_SVD(3:61)_ matrix to remove motion and noise, isolating the GV collapse event for

GV localization. We identified the first column of this matrix as the GV localization image comparable to the ground truth. This column is equivalent to the first B-mode frame with intact GVs before collapse, except that the clutter from transducer movement, Si particles, and noise has been filtered out. For this smaller dataset (frames 1 to 100), we did not apply grouped eigen-image difference indexing for changepoint detection, since in short sequences focused on identifying discrete transitions such as GV collapse, directly analyzing *E_t_* in (4) provides higher sensitivity. This enables more accurate detection of transient events without the potential loss of significant shift in the GV collapse signal that may occur with grouped difference averaging (*P_g_* in (7)).

## V. Conclusion

The eigen-image based thresholding approach, utilizing changepoint detection method for automatic segmentation of data, effectively detects ultrasound signals of point targets including MBs and GVs, separating from clutter and noise in a single-step process. This approach outperformed elbow point method and hard thresholding method by achieving the highest VD and SNR, resulting in clearer and detailed blood vessel visualization of the vessel-mimicking phantom and tumor in a tumor bearing mouse model. This approach also enables GV localization without pixel registration in a phantom by analyzing the GV collapse event while GVs are in apparent lateral motion.

